# Image-based leaf SPAD value and chlorophyll measurement using a mobile phone: enabling accessible and sustainable crop management

**DOI:** 10.1101/2025.07.08.663527

**Authors:** John Anderson, Kulaporn Boonyaves, Haruko Okamoto, Netiya Karaket, Wannisa Chuekong, Miguel Garvie, Napason Chabang, Daniel Osorio, Kanyaratt Supaibulwatana

**Author notes:** **Corresponding Authors** Daniel Osorio, Kanyaratt Supaibulwatana. **Competing Interests** Daniel Osorio and John Anderson are co-inventors on an international patent PCT/GB2006/000639, which covers the mobile phone imaging technology used in this study. Daniel Osorio, John Anderson, and Miguel Garvie are co-owners of the PhotoFolia mobile application, which was employed in the research. The other authors declare no competing interests.

## Abstract

This study evaluates a practical, low-cost solution for image-based leaf SPAD (Soil and Plant Analysis Development) value and chlorophyll content monitoring using a mobile phone. We compare laboratory assay and SPAD-502+ measurements with image-based estimates from a mobile phone app (PhotoFolia). Performance is tested for four commercial rice varieties grown in Thailand. Results show that the image-based method can predict SPAD values within ± 1.2 units Mean Absolute Error (MAE) and- chlorophyll concentrations within 7.2% Mean Absolute Percentage Error (MAPE) of laboratory results. Achieving a SPAD value error close to the industry standard of ±1 unit and a relative error of less than 10% in chlorophyll concentration estimation (compared to a laboratory method) demonstrates that an image-based approach using standard mobile phones can serve as an accessible, low-cost tool for on-farm chlorophyll monitoring, without the need for specialised equipment.

**Key Points / Highlights:** Novel low-cost approach for chlorophyll assay and SPAD-value measurement using standard mobile phone. Achieves accuracy comparable to commercial tools. Eliminates need for specialised sensors or laboratory equipment.

**Impact:** This study demonstrates that mobile phone-based image analysis can accurately estimate leaf SPAD and chlorophyll levels in rice under ambient lighting conditions, offering a low-cost, accessible tool for monitoring plant health.

## Introduction

Leaf chlorophyll concentration is a critical indicator of plant health. The strong correlation with soil nitrogen levels (Evans 1989) means that it provides valuable insight into nutrient status, stress, and overall productivity of plants. Monitoring chlorophyll levels enables farmers to optimize fertilizer use, improve crop yields, and detect early signs of environmental stress or nutrient deficiency (Peng et al. 1996). Effective chlorophyll monitoring in agricultural settings requires tools that offer ease of use, speed, accuracy and affordability.

Several methods are available for measuring chlorophyll concentration, ranging from colour-matching cards (IRRI 2005) to laboratory-based assays (Table 1). Each technique offers distinct advantages and limitations, making it suited to specific applications. While laboratory-based assays provide high precision, they are time-consuming, expensive, and require specialized expertise. Portable devices like the Minolta SPAD-502+ meter offer rapid field measurements from intact leaves but are prohibitively costly for many farmers. Furthermore, interpreting SPAD readings often requires calibration tailored to specific crops or conditions, adding complexity to their use (Markwell et al. 1995). While colour cards are low-cost their application benefits from experience, and results can suffer from human bias. Variations in illumination are a problem for image-based approaches, and mobile applications often integrate additional hardware to control illumination (Agarwal et al. 2021), which adds to their cost. Lastly, purely software-based mobile applications for chlorophyll assay offer the potential for widespread adoption due to their accessibility and ease of use, but they are regarded as the least precise (Kulig et al. 2024).

**Table 1.** Comparison of chlorophyll measurement techniques (Agarwal et al. 2021; Arnon 1949; IRRI 2005; Konica_Minolta_Sensing_Europe_B.V.; Petiole_LTD. 2017–2025: https://www.petiolepro.com/).

| Strategy | Speed | Ease of Use | Cost | Field Usability | Precision |
| --- | --- | --- | --- | --- | --- |
| Colour Cards<br>(IRRI 2005) | Moderate | Very Easy | Very Low | Excellent | Low to Moderate |
| Laboratory Assay<br>(Arnon 1949) | Slow | Challenging | High | Poor | Very High |
| Chlorophyll Meter<br>(Konica_Minolta_Sensing_Europe_B.V.) | Fast | Easy | Moderate | Excellent | High |
| Phone Apps, add hardware<br>(Agarwal et al. 2021) | Fast | Easy | Low | Excellent | Moderate |
| Phone Apps, software only<br>(Petiole_LTD. 2017–2025) | Fast | Extremely Easy | Very Low | Excellent | Low |

Most chlorophyll measurement techniques exploit the fact that chlorophyll absorbs light at specific wavelengths. Laboratory assays extract chlorophyll using solvents and infer chlorophyll concentration from measured absorption spectra. Portable meters like the SPAD-502+ estimate chlorophyll content non-destructively by comparing light transmitted through a leaf at a wavelength absorbed by chlorophyll relative to one that is not absorbed. Chlorophyll measurement can also take advantage of the light reflected by leaves. This is because reflected and transmitted light have similar spectral characteristics due to scattering by air-tissue interfaces in the mesophyll (Xu and Ye 2023). Remote sensing may employ hyperspectral cameras to record the spectral signatures of plants across large areas, enabling the mapping of chlorophyll and other pigments (Tucker 1979; Zimmermann et al. 2007). The rich structure of reflectance data has enabled the development of models of leaves that predict leaf properties from spectral images (Shi et al. 2024). The ability of algorithmic approaches to enhance the value of raw measurement data has led to software becoming increasingly important in chlorophyll monitoring. In addition, software-driven solutions, particularly those deployed via cloud services, are inherently scalable at low-cost compared to hardware systems, which depend on specialized supply chains and distribution networks. This creates an incentive to develop monitoring strategies that do not rely on specialised hardware. Notably, ordinary digital cameras produce an RGB signal by capturing reflected light through a colour filter array that separates incoming light into red, green, and blue wavelength bands corresponding roughly to the spectral sensitivities of cone receptors in the human eye. Each pixel in the resulting image contains intensity values for these three channels, usually encoded on a scale from 0 to 255, which together represent the colour of the scene. Because plant pigments like chlorophyll interact strongly with specific wavelengths - especially in the red and near-infrared regions - the spectral response captured in RGB images contains indirect but useful information about chlorophyll concentration. Recent advances in image processing and machine learning have made it feasible to treat chlorophyll measurement as an inference problem. The challenge has been to develop effective ways of mapping the camera signal onto chlorophyll concentration. Several studies have demonstrated consistent relationships between camera responses to leaves and chlorophyll concentration (Ali et al. 2012). However, many applications require careful control of the way that the leaf is illuminated, either using a known light source, such as the camera flash, in conjunction with a specialised hardware enclosure (Agarwal et al. 2021) or relying on contact images (Ibrahim et al. 2021). These also require a controlled light source shining through a leaf placed over the camera aperture to ensure a consistent camera response. Despite demonstrating strong correlations between the camera response and chlorophyll concentration, the need for additional hardware limits the reach of these applications.

Purely software-based approaches do exist that attempt to quantify the greenness of leaves and relate this to chlorophyll concentration, but they do not explicitly compensate for lighting conditions or camera hardware and often rely on the use of controlled lighting conditions or a reference card to white balance the image and normalise the brightness. The Dark Green Colour Index (DGCI) first converts the camera RGB value into the HSV colorimetric space. The HSV colour space represents colours using hue (H) angle of colour on the colour wheel, saturation (S) the intensity or purity of the colour, and value (V) the brightness, offering an intuitive alternative to RGB that aligns closely with human perception. Pixels that fall within a particular hue angular range are associated with green leaf colours. These pixels are then classified based on their saturation and brightness to give a measure of greenness and an overall measure is derived for a given leaf area. The Green Leaf Index (GLI) is a similar measure, which involves calculating a ratio from the RGB signal that supresses red and blue signals while emphasising green. Healthy leaves containing more chlorophyll yield higher GLI and DGCI values. However, these measures are primarily used for relative comparison rather than for recovering absolute chlorophyll levels.

Despite the growing availability of such indices and related mobile applications, there remains a lack of robust validation for image-based algorithmic methods when benchmarked against conventional specialist hardware solutions such as SPAD meters or spectrophotometers. This may reflect the relatively low barrier to entry for app development, where the low cost of deployment often leads to lower user expectations for accuracy or consistency. Consequently, developers may have little incentive to conduct rigorous performance testing or calibration across different lighting conditions or device types.

There is also justification for scepticism about whether subtle variations in leaf colour can be objectively measured and reliably linked to chlorophyll concentration using images captured by standard consumer-grade cameras. The RGB signal in most digital cameras compresses the visible spectrum into three broad spectral bands (red, green, and blue) which inherently limits spectral resolution. Moreover, this signal is influenced by external factors such as ambient illumination, camera sensor characteristics, and image processing algorithms, making consistent and objective quantification of biophysical properties like chlorophyll content challenging. As a result, while DGCI and GLI offer accessible and non-destructive proxies for assessing plant health, their reliability and reproducibility remain constrained by the limitations of the imaging systems on which they rely.

To address this knowledge gap, we compare three types of chlorophyll measurement: a standard laboratory-based assay (spectrophotometric measurement), a SPAD-502+ meter, and estimates derived from mobile phone images of leaves obtained under ambient conditions (PhotoFolia app).

## Materials & Methods

### Rice growth protocol

Seeds of four Thai rice, *Oryza sativa* ssp., Indica cultivars, Khao Dawk Mali 105 (KDML), RD6, RD47, and Pathum Thani 1 (PPT1) were obtained from Pathum Thani Rice Research Centre, Thailand (Jiamtaea et al. 2017). Seeds were surface sterilised and were imbibed and kept at 30°C for 5 days to induce germination. The vigorous germinated seeds were transferred to a hydroponic-based microgreen system (AgrowLab® Co.Ltd., Thailand) supplemented with Yoshida solution (Yoshida 1976) and incubated under a 16 h d^-1^ photoperiod at 80 mmol photons m^-2^ s^-1^ of LED lighting (445, 554 nm) for three weeks. Seedlings of each of the four cultivars with uniform growth were then individually transferred to vermiculite-based substrate culture supplemented with nitrogen levels of 0.26 mM NH_4_NO_3_ or 2.056 mM NH_4_NO_3_ of modified Yoshida formulation, and placed in a Mahidol University greenhouse (Phayathai campus) with average temperature at 33.8+6.4 °C. and 17,409+1,881 lux illumination (recorded by the Data logger model UA-002-64; HOBO Pendant® Temperature/Light 64K Data Logger). Assuming natural daylight conditions, this corresponds to a photosynthetic photon flux density of approximately 322 ± 35 µmol m^−2^ s^−1^, using a standard conversion factor of 0.0185 µmol m^−2^ s^−1^ per lux for sunlight (Thimijan and Heins 1983). The experiments were set up in the greenhouse on August 14, 2024, and all parameters recorded after 21 days of cultivation on September 4, 2024.

### Leaf measurement protocols

For each cultivar–condition combination, three independent biological replicates were established, resulting in a total of 24 individual plants (4 cultivars × 2 conditions × 3 biological replicates). At harvest, the third leaf (counting from the shoot apical meristem) was identified on each plant. From this leaf, the middle two-thirds of its length was excised to avoid highly variable regions near the base and tip (Yuan et al. 2016). This mid-section was then divided into five approximately 3-cm segments, yielding five subsamples per plant. These segments served as pseudo-replicates (i.e., spatial subsamples) to account for within-leaf heterogeneity and to provide sufficient material for robust comparison of the four different measurement techniques described below.

Leaf images were first collected using the PhotoFolia app and colour standard. The PhotoFolia app is available for both Android and iOS operating systems (https://www.colourworker.com/apps). Three models of iPhone were used in this study (14 Pro, 14 Pro Max & 15 Pro Max). The PhotoFolia app was downloaded from the Apple (iOS) store using the QR code printed on the colour standard. After registering (Google/Apple/Facebook) the class of reference reflectance spectra (rice variety/condition) was selected before taking a photograph of the colour standard together with a rice leaf (Fig. 1a).

**Fig. 1.**
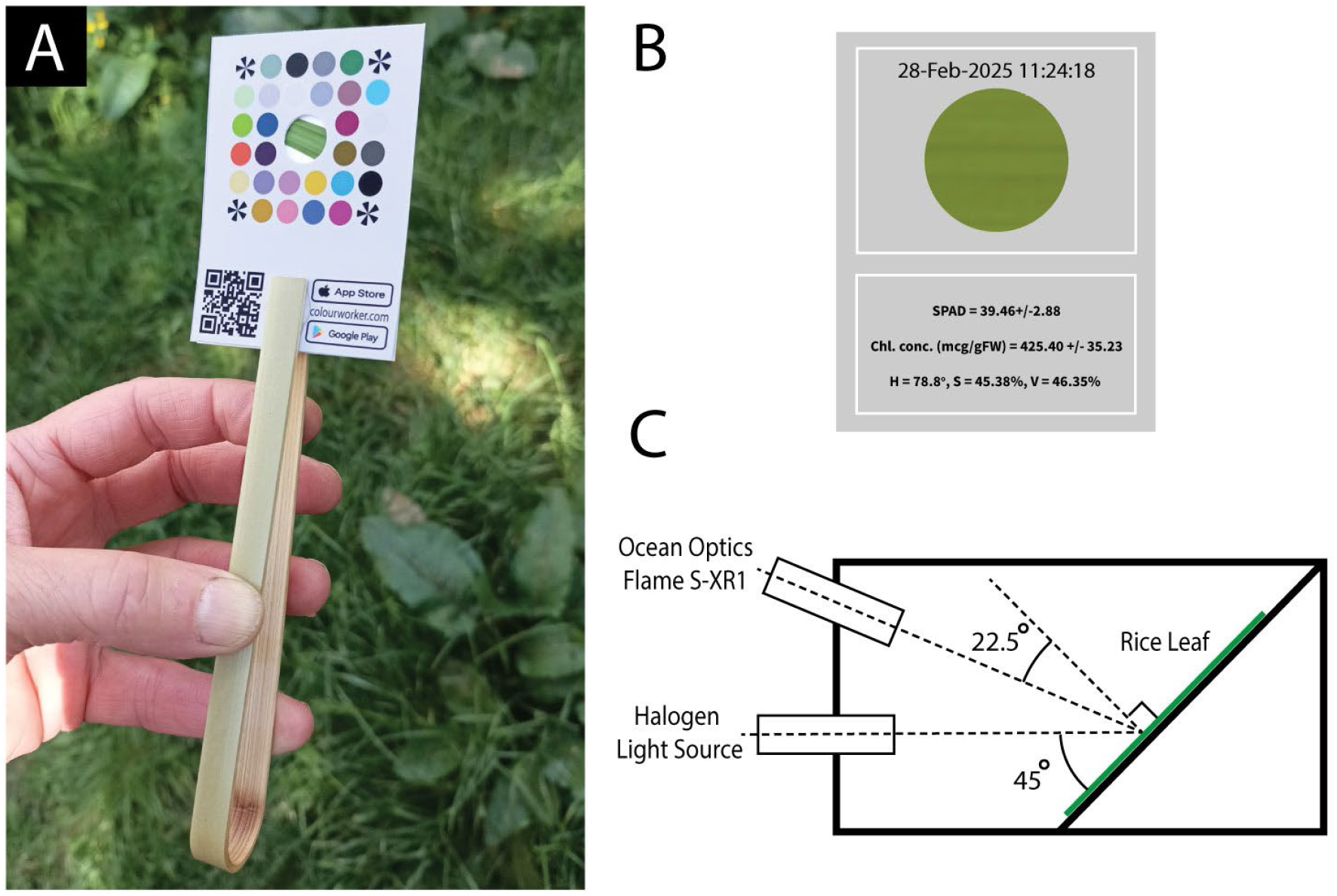
A) Colour standard tweezers enable a leaf segment to be gripped flush with the surface of the standard and aligned with the sample aperture. The user takes a photograph of the colour standard from within the PhotoFolia app taking care to include the radial corner location motifs. B) PhotoFolia results screen displays the leaf sample region together with SPAD value, chlorophyll concentration and HSV colorimetric coordinates. C) Reflectance Spectrum measurement schematic. The leaf is mounted at 45° inside a light tight matt black painted aluminium housing. Illumination is via a microscope fibre optic halogen light source at 22.5° to leaf normal. Spectral reflectance measured at 45° to leaf normal via fibre optic focused with collimating lens and Ocean Optics Flame S-XR1 spectroradiometer using OceanView software.

The colour standard is mounted on a pair of bamboo forceps enabling a leaf section to be gripped, pressed flush and aligned with a 10mm diameter circular aperture in the centre of the standard (Fig. 1a). Under uniform illumination ensures that both the leaf and the standard receive the same light. The leaf segment was gripped so that the central portion of the leaf aligned with the sample aperture.

The acquired image is uploaded to a cloud-server (AWS) running the Matlab Production Server (Mathworks) where image-processing takes place before results are returned to the mobile phone client for display. An example of the PhotoFolia app results screen (Fig. 1b) displays the sample region together with estimates of SPAD value, chlorophyll concentration and (HSV) colorimetric coordinates. Historical image data and estimates can also be browsed and downloaded from a web application that is accessible via the account used when registering the PhotoFolia app (https://www.colourworker.com/webapps).

A single SPAD reading was taken using a Minolta SPAD-502+ chlorophyll meter (Konica Minolta) from the central region of the leaf segment, ensuring the measurement area overlapped as closely as possible with the image-based and reflectance analyses. The meter was calibrated according to the manufacturer’s instructions prior to use.

To measure leaf reflectance spectra, the leaf section was mounted in a custom-built aluminium light tight enclosure (85 x 60 x 60mm) allowing the adaxial leaf surface to be diffusely illuminated by a Halogen fibre-optic illuminator oriented at 45 ° to the surface normal (Fig. 1c). A 200µm UV-VIS fibre optic was oriented at 22.5° to the leaf surface normal to minimise specular reflection. The tip of the fibre optic was focused via a collimating lens to sample from an elliptical region (∼5mm minor axis) on the leaf surface. The spectral distribution of reflected light was measured using an Ocean Optics Flame S-XR1 Spectroradiometer with instrument dark spectrum subtracted and calibrated against a PTFE white standard.

Following optical measurements, the same segment of leaf was weighed (+/-0.1 mg) and snap frozen in liquid nitrogen for chlorophyll extraction. Chlorophyll was extracted by grinding the tissues using a homogeniser in 80% acetone and the absorption maxima of chlorophyll a (662 nm) and b (644 nm) were measured by a spectrophotometer (Thermo-scientific Genesys 10s uv-vis spectrophotometer, METTLER TOLEDO^©^, USA). Extractions were made under normal lab lighting conditions though samples were placed in lidded ice-buckets during the extraction steps to minimise degradation of chlorophyll. Total chlorophyll per gram fresh weight of each tissue was calculated by the formula published by Lichtenthaler (Lichtenthaler 1987).

### Image-based estimates of chlorophyll concentration and SPAD value using PhotoFolia

Recovery of objective information about leaves from ordinary camera images presents two challenges. First, the camera RGB signal (V) depends on the spectral composition of the illuminant (I), the spectral reflectance of the leaf (R) and the spectral sensitivity of the camera sensor (S).

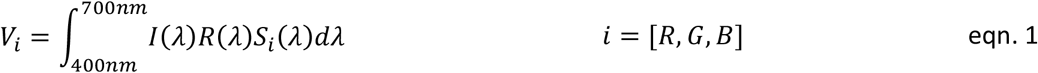

This relationship means a leaf photographed with a different make/model of camera or under different lighting conditions may produce a different camera signal. It is therefore necessary to compensate the camera response to make it invariant to the illuminant and to camera spectral sensitivity while preserving information about changes in leaf spectral reflectance. The second challenge is that although changes in chlorophyll concentration influence the spectral reflectance of the leaf, understanding how the camera responds to these changes is necessary to infer chlorophyll levels from the camera signal. The PhotoFolia app address both of these challenges (Fig. 2) to enable an objective measure of chlorophyll content (or SPAD value) to be recovered from leaf images.

**Fig. 2.**
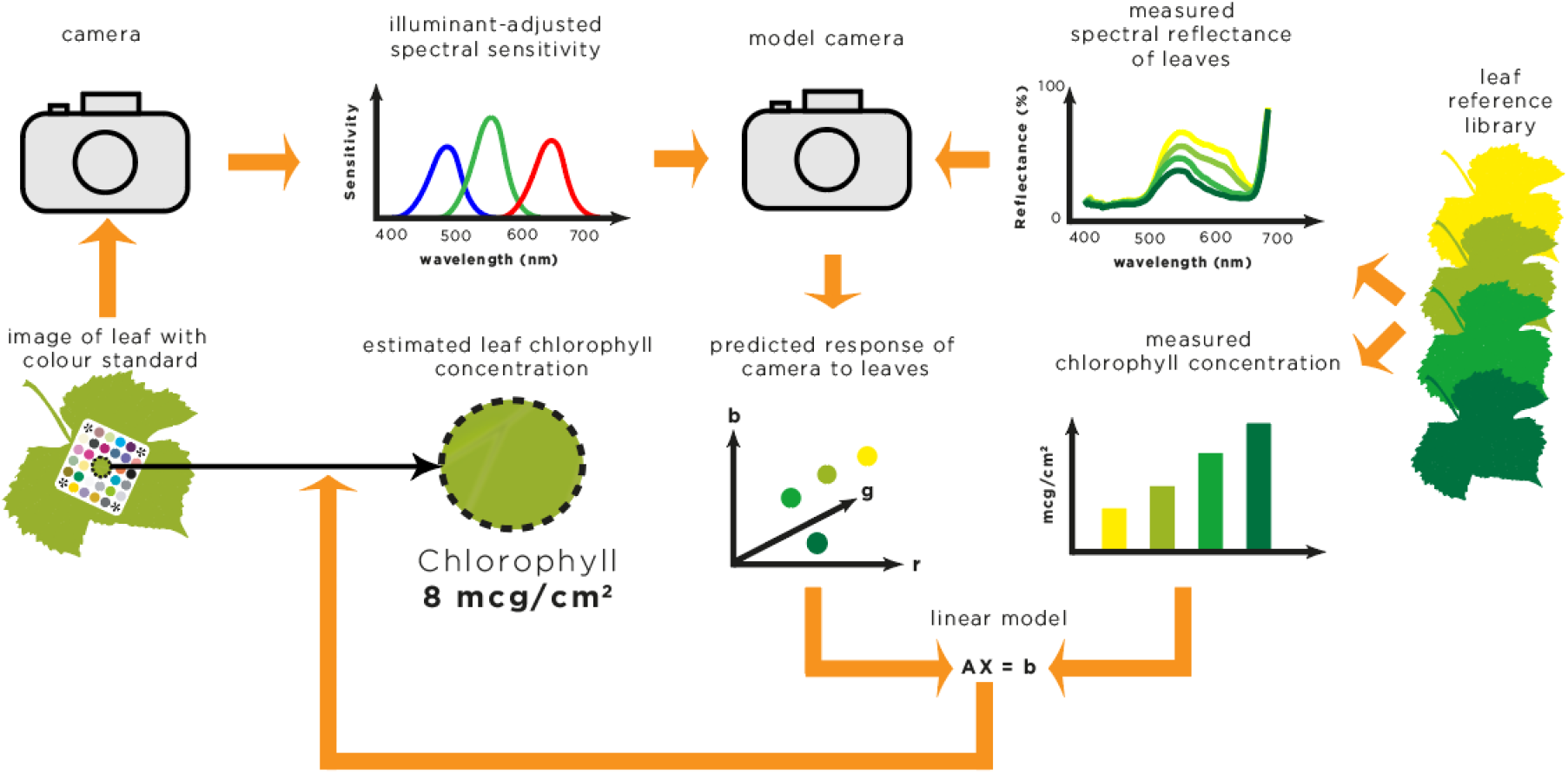
Schematic of algorithm for chlorophyll measurement implemented by the PhotoFolia app. Left side: a real image of a leaf containing a colour standard with coloured patches of known spectral reflectance is used to estimate a model of the spectral sensitivities of the camera’s red, green and blue sensors that also takes account of the scene illumination. Centre: the model camera is used to predict the response of the real camera to a set of reference reflectance spectra of leaves containing different levels of chlorophyll (right). Bottom-right: measured leaf chlorophyll concentrations are regressed on to the estimated model camera response to the reference reflectance spectra. This linear model is then used to predict the chlorophyll concentration from real image RGB values recovered from the leaf (bottom-left). The same procedure is applied when using PhotoFolia to estimate SPAD value. In this case measured SPAD values associated with leaf spectral reflectance curves are used during the regression step.

To compensate for variations in scene illumination and make/model of camera, PhotoFolia employs a colour standard comprised of coloured patches of known spectral reflectance. The camera RGB signals associated with the patches recovered from an image of the colour standard are used to estimate a model of the spectral encoding characteristics of each of the camera’s colour channels that also takes account of the scene illumination. This is achieved by a constrained least-squares optimisation (Finlayson et al. 1998). The resulting camera model can then predict how the real camera will respond to leaf reflectance spectra.

Given a sample of leaf reflectance spectra that are paired with SPAD or chlorophyll assay measurements it is possible to train a model to predict chlorophyll concentration or SPAD value from the estimated camera response to the leaf reflectance spectra. The resulting model can then predict SPAD value or chlorophyll levels from RGB values recovered from actual leaf images. If the variation in the spectral reflectance of the subject material can be described using relatively few degrees of freedom it is possible to recover accurate estimates from the camera signal (Chiao et al. 2000), and makes (near) optimal use of camera RGB signals to estimate spectral reflectance - or any quantity that is correlated with spectral reflectance.

A detailed evaluation of the accuracy of camera models is beyond the scope of this study. However, the adopted quadratic programming approach (Finlayson et al. 1998) is an effective strategy for finding robust, accurate and reproducible solutions to inherently ill-conditioned optimisation problems.

### The relationship between camera response and chlorophyll concentration and SPAD value

The relationship between chlorophyll concentration, SPAD value, and light interaction with leaves has been extensively studied (Gausman 1985; Uddling et al. 2007). The primary aim here is to select an appropriate model for relating RGB signals derived from leaf images to SPAD values and chlorophyll concentration, which requires brief consideration of the optical processes governing light propagation in leaf tissue.

The Beer–Lambert law describes light attenuation in a homogeneous, non-scattering medium, where absorbance A is proportional to the concentration c of the absorbing species, its molar extinction coefficient ε, and the optical path length l:

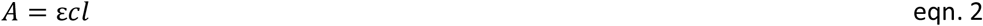

Absorbance is defined as:

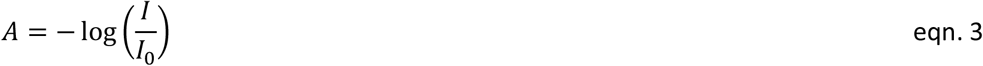

where I_0_ is the incident light intensity and I is the transmitted intensity. This formulation arises from the assumption that the fractional loss of light intensity per unit path length is proportional to the current intensity. Combining Eqs. 2 and 3 yields:

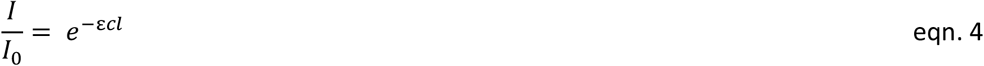

indicating that transmitted light decreases exponentially with increasing absorber concentration and path length. However, leaves are not homogenous structures containing a uniform distribution of chlorophyll. They are heterogenous cellular structures that scatter light in complex ways meaning that the Beer–Lambert law does not strictly apply (Gausman 1985).

The Kubelka-Munk theory (Kubelka and Munk 1931) provides a foundational framework for modelling light transport in diffusely scattering media such as plant leaves (Jacquemoud and Baret 1990; Zhang et al. 2017). Unlike simple exponential attenuation - typical of non-scattering media - light in strongly absorbing, highly scattering tissues like leaves exhibits more complex behaviour, often approximating a reciprocal decay in intensity under certain conditions. However, empirical studies have shown that neither pure exponential nor reciprocal decay adequately captures the nuanced relationship between chlorophyll concentration and the intensity of reflected or transmitted light (Gitelson and Merzlyak 1994; Thomas and Gausman 1977). This complexity arises because, in a leaf, incident photons undergo multiple scattering events while being progressively absorbed by chlorophyll. One important consequence of this for our study is that this results in similar spectral signatures in both reflected and transmitted light (Xu and Ye 2023).

For practical purposes, over a relatively small dynamic range 1/C and exp(-C) have a similar shape and while theoretically they reflect entirely different processes, within the bounds of statistical variations they capture a similar characteristic, that of a non-linear decrease in the intensity of reflected light as chlorophyll concentration increases. Though it is clearly an oversimplification to write:

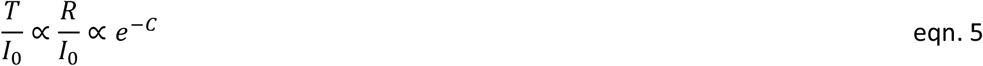

this is not intended to imply precise compliance with Beer–Lambert physics but rather reflects the dominant role of chlorophyll absorption in attenuating light within a complex scattering medium (Jacquemoud and Baret 1990).

The Minolta SPAD-502 meter aligns with this assumption and leverages the strong absorption of chlorophyll at 650 nm and uses a reference wavelength at 940 nm (where chlorophyll absorption is negligible) to generate a stable chlorophyll index that is approximately linearly related to chlorophyll concentration. Although the exact internal calibration is proprietary, published formulations (Uddling et al. 2007) express SPAD as:

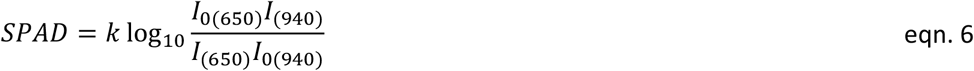

This dual-wavelength approach partially compensates for variations in leaf thickness and internal structure, as the 940 nm NIR signal compensates for non-pigment-related attenuation (e.g., scattering, path length effects). Nevertheless, residual dependence on leaf anatomy can persist, particularly across species with divergent leaf structures (Marenco et al. 2009). In our study, rice leaves exhibited relatively uniform thickness, so we expect structural variability to be minimal, and SPAD values to primarily reflect differences in chlorophyll content per unit area.

Although Eqn. 6 implies a linear relationship between SPAD and chlorophyll concentration, empirical studies consistently report a non-linear (saturating) relationship at high chlorophyll levels (Markwell et al. 1995; Monje and Bugbee 1992). This deviation arises from the breakdown of underlying assumptions in real leaf tissue.

Our focus, however, is on the relationship between camera-derived RGB signals and SPAD/chlorophyll measurements. Modern CMOS sensors exhibit a linear response to photon flux (Eqn. 1), meaning the digital response (V) is proportional to reflected light intensity (R). Given that reflectance in the red band decreases with chlorophyll content, the camera response is expected to decline approximately exponentially with increasing chlorophyll content and SPAD value:

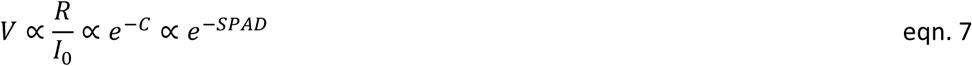

Consequently, the logarithm of the camera response should exhibit an approximately linear, inverse relationship with SPAD and chlorophyll concentration:

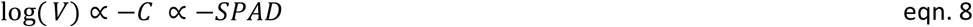

This suggests that a linear model regressing SPAD (or chlorophyll) against the log-transformed camera signal (V) is a reasonable starting point. However, because the exponential approximation may not hold universally, we empirically compared linear models fitted to both raw and log-transformed camera responses.

### Choice of reference reflectance spectra when making estimates using PhotoFolia

PhotoFolia requires reference reflectance spectra to train the model for predicting SPAD values and chlorophyll concentrations from leaf images (Fig. 2). This step is critical because the model must account for the spectral characteristics of light reflected by the leaf. However, the appropriate level of specificity when selecting reference spectra requires consideration. If all leaves exhibited identical changes in spectral reflectance as a function of chlorophyll concentration, a “generic” set of reference spectra would suffice. However, leaf spectral reflectance is influenced by multiple factors beyond chlorophyll concentration, including structural features such as ridges, hairs, and surface roughness, and secondary pigments including carotenoids and anthocyanins. Furthermore, there is no universal mapping between hardware-based SPAD measurements and leaf chlorophyll content (Uddling et al. 2007) so a single set of reference spectra may not adequately capture species-specific variations in the relationship between spectral reflectance and chlorophyll concentration.

To evaluate the choice of reference spectra on predictive error this study compared three types of reference library:

1. A generic library combining spectral reflectance measurements across all rice varieties and experimental conditions.
2. Variety-specific libraries, which included reflectance spectra from both control and treatment groups for each rice variety.
3. Variety-specific and group-specific libraries, which separated reflectance spectra by both rice variety and experimental condition.

In each case, the appropriate variety- and/or group-specific reference spectra were applied when estimating SPAD values and chlorophyll concentrations for a given rice variety and experimental condition.

### PhotoFolia (training and testing)

PhotoFolia uses the reference spectra to calibrate a predictive model for estimating SPAD values and chlorophyll concentrations from image RGB values. To optimise the use of a relatively small dataset, a leave-one-out cross-validation (LOOCV) strategy was adopted for evaluating model performance. This approach maximises use of available data during training while rigorously excluding any information pertaining to the spectral reflectance, SPAD values, or chlorophyll concentrations of the test sample, thereby maintaining the integrity of the validation procedure.

The predictive model for estimating SPAD values or chlorophyll concentrations from RGB signals takes the linear form b = A X, where b is an n×1 vector of observed SPAD values or chlorophyll concentrations from rice leaves, A is an n×3 matrix containing either raw RGB values or their logarithmic transformation (log(RGB)), and X is the 3×1 vector of unknown model coefficients to be estimated. Due to strong collinearity among the camera’s RGB responses—arising from the broad and overlapping influence of chlorophyll on leaf reflectance spectra—ordinary least-squares estimation can yield unstable or unreliable coefficient estimates. To address this, ridge regression was employed as a regularization technique, which introduces a penalty on the size of the coefficients to improve numerical stability and generalization. The strength of this penalty is controlled by the regularization parameter λ. To select an optimal value for λ, we used a grid search: a systematic procedure that evaluates model performance across a predefined set of candidate λ values. For each candidate, model accuracy was assessed using 3-fold cross-validation, and the λ yielding the best average performance was chosen.

Three metrics were used to evaluate the accuracy of estimated SPAD values and chlorophyll concentrations against measured values. The Mean Absolute Error (MAE) measures the average absolute difference between estimated and measured values, providing a straightforward assessment of performance that is independent of the magnitude of the values. MAE provides an intuitive sense of the typical error size in the same units as the data, making it easy to interpret. Additionally, the Mean Absolute Percentage Error (MAPE) assesses accuracy relative to the magnitude of the measurements and is a useful measure of utility, in particular identifying potential bias in estimates at low values. Finally, R-squared (R^2^) values were calculated as an indication of the proportion of variance in the measured values that is explained by the estimated values. This measure is useful for assessing the overall goodness-of-fit of the model and reveals whether the estimation approach is effectively taking advantage of structure in the training data. Bland-Altman plots were also used to reveal obvious structure in the errors as a function of estimation value.

## Results

### Camera response to leaf reflectance spectra vs measured SPAD values and chlorophyll concentration

To comprehensively examine the relationship between the camera response, measured SPAD value and chlorophyll concentration, it is necessary to ensure controlled illumination during imaging and precise tracking of a defined spatial region of the leaf during imaging, SPAD measurement, and chlorophyll assay. A standardised protocol of this kind was not applied in this study. However, it is possible to gain insight into the relationship between the camera response and measured SPAD value and chlorophyll concentration by assuming that the camera is linear. This is a reasonable assumption given that CMOS sensors exhibit a linear photoelectric response and are universal in mobile devices. Using eqn. 1 the response of the camera can be calculated from an estimate of the illuminant-adjusted spectral sensitivities and the measured reflectance spectra of different leaves. The calculated camera responses will approximate the real camera response to the leaves, and this can be compared to measured SPAD values and chlorophyll concentrations from the same leaves. This relationship can help to inform the most appropriate choice of model for making predictions from the real camera RGB values.

The camera response for a given channel is modelled by the standard spectral integration equation (eqn. 1). Using this model, we first estimated the spectral sensitivities of a representative camera from its responses to the colour standard. We then applied Equation 1 to compute the predicted green channel response (V_G_) for each measured leaf reflectance spectrum. These predicted responses are plotted against measured SPAD values in Figure 3 and against laboratory chlorophyll concentrations in Figure 4.

**Fig. 3.**
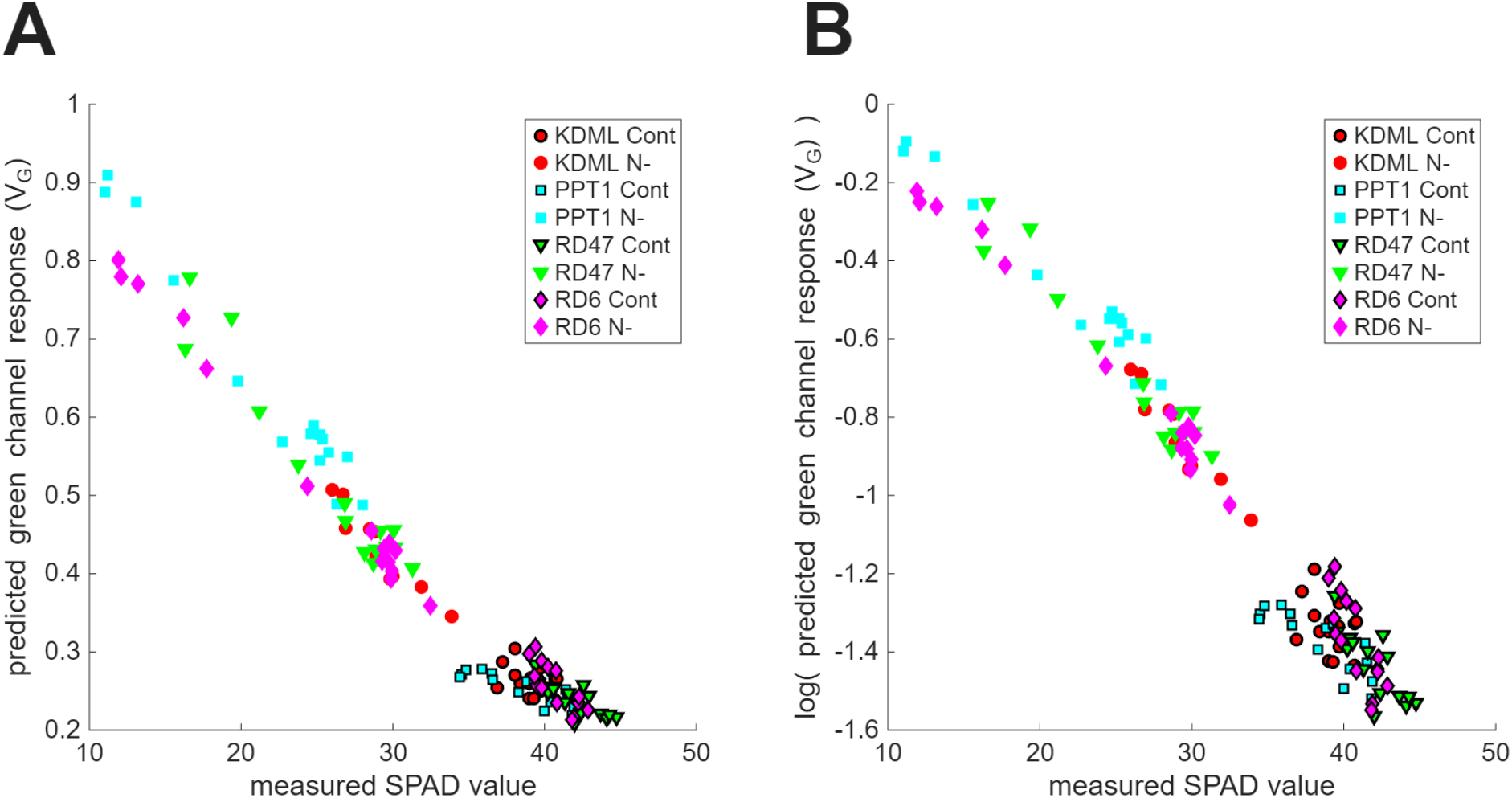
Comparison of predicted green channel model camera (linear) response (V_G_) to reflectance spectra obtained from seven-week-old leaves of KDML, PPT1, RD47, and RD6 rice varieties grown under two nitrogen concentrations Cont (2.056 mM), and N- (0.26 mM) with SPAD value measurements from same leaves. (A) camera response is inversely proportional to measured SPAD value, with an asymptote apparent at high SPAD values. (B) log transformation of the camera response linearizes the relationship at high SPAD values but results in a loss of linearity at low SPAD values.

**Fig. 4.**
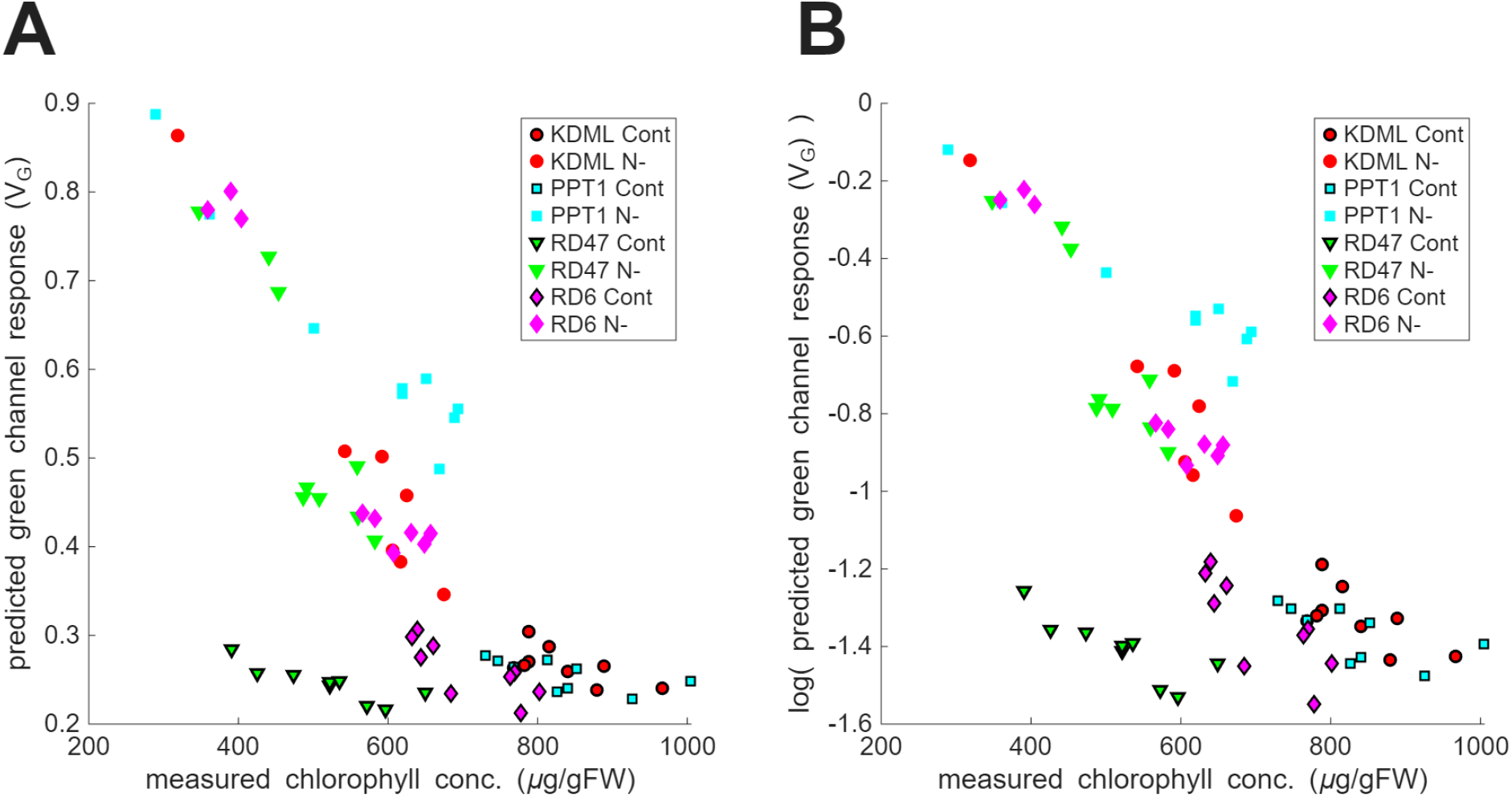
Comparison of predicted green channel model camera (linear) response (V_G_) to reflectance spectra obtained from seven-week-old leaves of KDML, PPT1, RD47, and RD6 rice varieties grown under two nitrogen concentrations Cont (2.056 mM), and N-(0.26 mM) with chlorophyll concentration measurements (µg/gFW) from the same leaves. (A) There is an inverse relationship, between the camera response and measured chlorophyll concentration, however, this is not generally consistent and depends on the rice variety and growth condition. There is some suggestion of an asymptote of the camera response in control groups at high chlorophyll concentrations although the inverse relation still holds, indicating that the camera is responding to changes in chlorophyll concentration. (B) Log transformation of the camera response largely preserves the inverse linear relationship and also slightly reduces the asymptote of the camera response at high chlorophyll concentrations observed in the control groups. However, there is some indication of the introduction of a slight asymptote at low chlorophyll concentrations.

Although all three RGB channels were computed, only the green channel is shown because the channel responses are highly correlated. This correlation arises because leaf reflectance varies smoothly across a broad wavelength range while camera colour filters have overlapping, broadband sensitivities - making the channels convey largely redundant information for this application.

In Fig. 3 the untransformed green channel response (Fig. 3A) and the logarithm of the response (Fig. 3B) are plotted against SPAD value. The observed negative correlation between the SPAD value and the camera response is expected because the more light that is absorbed by chlorophyll in the leaf (higher SPAD value) the less light will be reflected (lower reflectance). By definition (Eq. 1), the camera response is linearly proportional to spectral reflectance. Because SPAD values relate logarithmically to chlorophyll content—and thus to reflectance—we expect a logarithmic relationship between SPAD and camera response. This is only weakly evident in the data as a slight asymptotic flattening of the camera response at high SPAD values (Fig. 3A). Applying a logarithmic transformation to the camera response improves linearity in this high-SPAD range (Fig. 3B); however, it compromises the otherwise linear response observed for less green leaves (i.e., at lower SPAD values). The benefit of logarithmically transforming the camera response was investigated by fitting linear models to predict SPAD values given the untransformed and log-transformed camera data. Log transforming the camera response resulted in an increase in both the MAE and the MAPE from 0.78 SPAD units and 2.65% to 0.84 SPAD units & 2.80% respectively, which favours the linear model to predict SPAD value from the camera signal for real image data.

As expected, there is a negative correlation between chlorophyll concentration and the predicted green channel response (Fig. 4) but the relationship between concentration and camera response depends on both rice variety and experimental treatment. Once again, there is an indication that the log transformation of the camera signal linearises the relationship at higher chlorophyll concentrations, although the control groups (Fig. 4A, black edged symbols) display a reasonable linear relation, albeit over a smaller range of green channel excitation. In addition, as for the SPAD/camera response, logarithmic transformation introduces a slight asymptote at low chlorophyll concentrations (Fig. 4B). Instead of fitting a single model to the relationship between the predicted camera response and the chlorophyll concentrations, the mean MAE and MAPE values were calculated from models fitted to variety and condition specific data for untransformed and log transformed camera responses. As for the SPAD data, logarithmically transforming the camera response resulted in an increase in the MAE and MAPE from 26.08 ug/g FW and 4.31% to 27.03 ug/g FW & 4.37% respectively. This is a minor difference, but, as for SPAD estimates, the fact that lower MAE and MAPE were associated with using untransformed camera data favoured applying a purely linear model to predict chlorophyll concentration from the camera signal.

Plotting measured SPAD value against measured chlorophyll concentration (Fig. 5) reveals the inconsistent relationship between these two quantities. While rice variety and condition separately display reasonable linear relationships indicated by linear fits and 95% confidence intervals on the slope parameter, there are noticeable variations between groups. In particular, the control and treatment groups associated with the rice variety RD47 are not continuous with one another. Similar chlorophyll concentrations in both groups are associated with quite different SPAD values. This study is not intended to provide a detailed assessment of the relationship between measured SPAD value and directly measured chlorophyll concentration, but these results imply that while a single generic model relating the camera response to SPAD value might be a valid choice, this is unlikely to be effective for predicting leaf chlorophyll content. This is because the variety- and condition-specific changes in chlorophyll concentration as a function of SPAD values differs substantially from the average change.

**Fig. 5.**
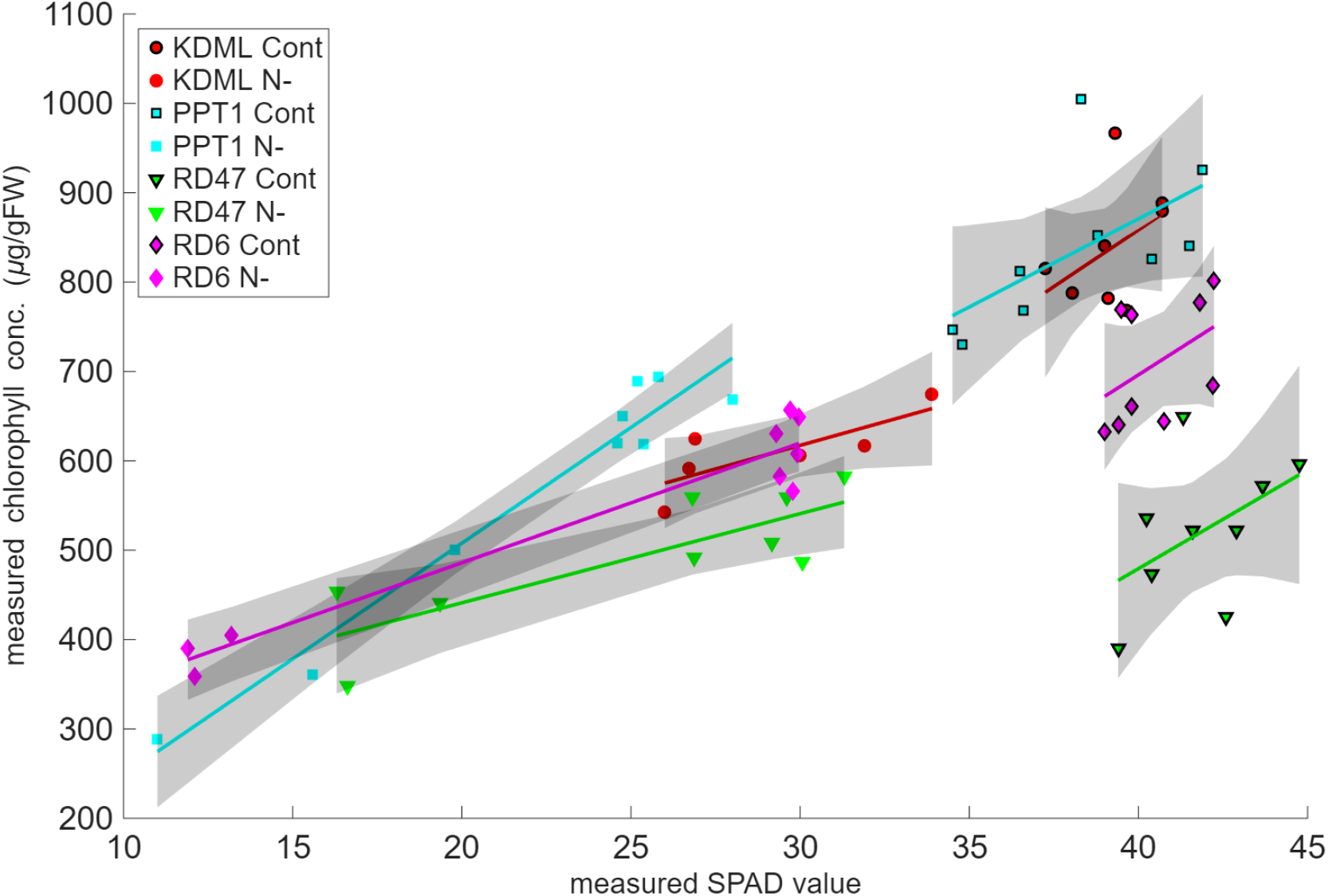
Comparison of measured chlorophyll concentration (µg/gFW) and measured SPAD values from the same seven-week-old rice leaves of KDML, PPT1, RD47, and RD6 rice varieties grown under two nitrogen concentrations Cont (2.056 mM), and N- (0.26 mM). Linear fits together with 95% confidence intervals on the slope parameter are indicated in grey. All rice varieties and conditions display a positive linear correlation. The correlation is weaker in the control groups partly as a result of the reduced measurement range. There is a strong indication of variety and condition specific variations in the relationship (see text).

### Choice of reference reflectance spectra when making estimates using PhotoFolia

To evaluate how the choice of reference spectra affects PhotoFolia estimates, we compared SPAD values and chlorophyll concentrations obtained from real camera images of leaves to their corresponding measured values, while varying the specificity of the reference spectra used (Fig. 6). Lines of equality are plotted in black.

**Fig. 6.**
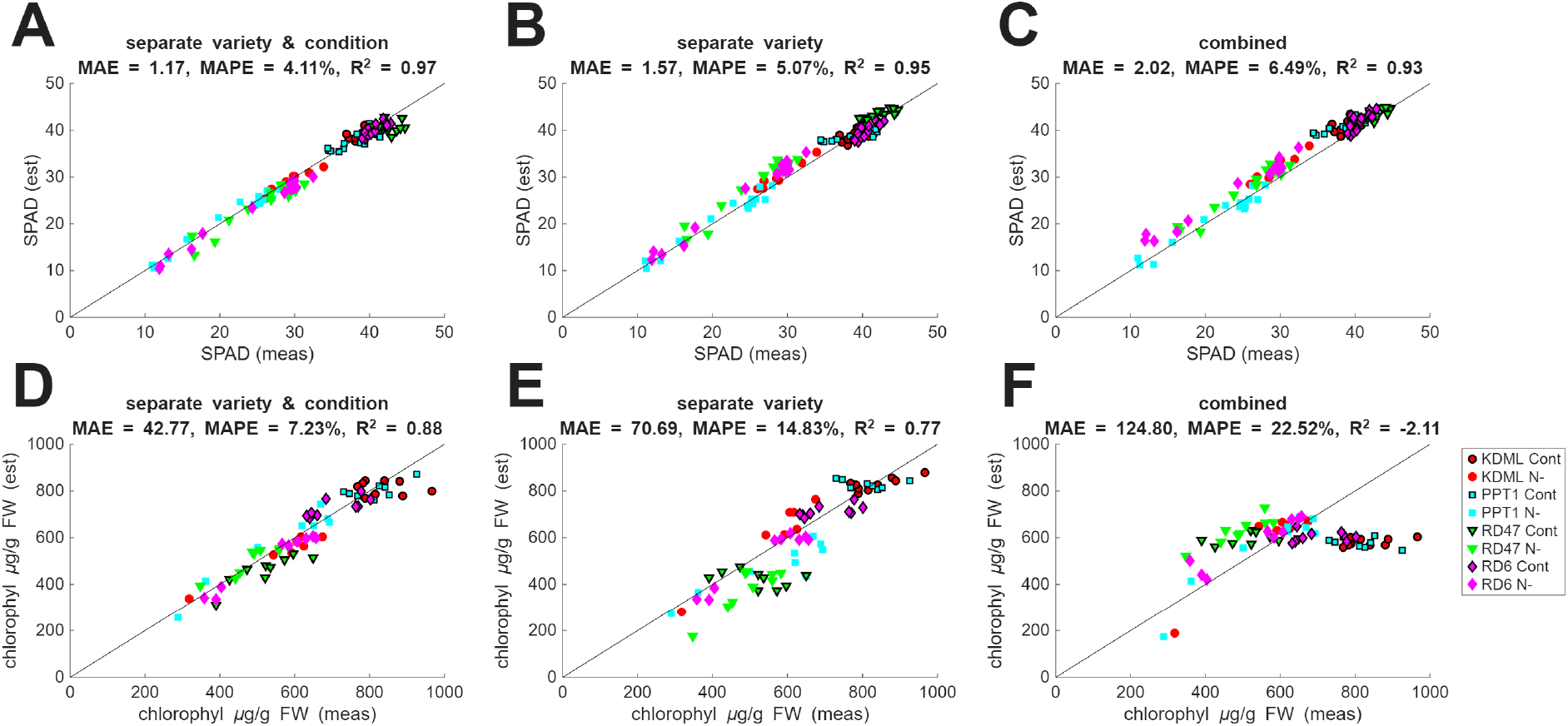
Comparison of measured SPAD value (A, B, C) and chlorophyll concentration (D, E, F) with image-based estimates made using PhotoFolia app showing the impact of varying the specificity of leaf reference reflectance spectra (left to right). Seven-week-old leaves of KDML, PPT1, RD47, and RD6 rice varieties grown under two nitrogen concentrations Cont (2.056 mM), and N-(0.26 mM) were tested. (A, D) separate variety, control and treatment specific reference spectra are used. (B, E) control and treatment reflectance spectra are merged into variety-specific groups. (C, F) all reference spectra are merged into a single group. Decreasing specificity (left to right) has a small impact on SPAD value estimation accuracy. MAE increases from 1.17 to 2.02 SPAD units and estimates remain strongly correlated with measurements. In contrast, the accuracy of chlorophyll estimates decreases substantially as reference reflectance spectra specificity reduces from a MAPE of 7.23% to 22.5%. There is a complete loss of correlation between estimates and measurements when utilising the combined reference reflectance spectra. This emphasises the value of prior knowledge regarding plant variety and growth conditions when inferring physical properties of leaves from optical measurements.

The scatterplots (Fig. 6A, B, C) compare rice leaf SPAD values measured using a Minolta 502+ SPAD meter and SPAD values estimated from images of the same leaves using PhotoFolia. The scatterplot 6A results from using variety- and condition-specific reference reflectance spectra (separate treatment and controls corresponding to the most specific configuration) when making estimates. The scatterplot 6B makes use of variety-specific reference spectra (combining control and treatment spectra for each rice variety) and 6C combines all reference spectra across rice variety and condition (least specific configuration).

As reference spectra specificity decreases (left to right) MAE and MAPE increase while R^2^ decreases but the effect is quite small. The relative error (MAPE) which is a useful summary of overall performance increases from 4.1% to 6.6% when moving from using the most specific to least specific reference spectra.

For chlorophyll estimates (Fig. 6D, E, F) the choice of reference spectra is more significant than for SPAD value. The relative error increases from 7.2% when using variety- and condition-specific reference spectra to 22.5% when applying combined reference spectra. Using combined reference spectra also result in a negative R^2^ value and there is no visual indication of a correlation between measured and estimated values.

Bland-Altman plots (Fig. 7) compare the difference between measured and estimated values as a function of average value and align to plots on Fig. 6. In the case of SPAD value there is a small but significant bias (p<0.01 t-test) resulting in a slightly higher estimated SPAD value compared to the measured SPAD value. This bias is lowest when applying the most specific reference spectra (0.72 SPAD units, Fig. 7A) and highest for the combined reference (1.79 SPAD units, Fig. 7C). The limits of agreement (LoA) increase as reference library specificity decreases (left to right). However, even when applying the combined reference, most estimated SPAD values lie within +/- 3 SPAD units of the measured values (Fig. 7C). There is some indication that for estimates associated with leaves belonging to the control groups, errors correlate with SPAD value, however, these errors still display relatively little bias.

**Fig. 7.**
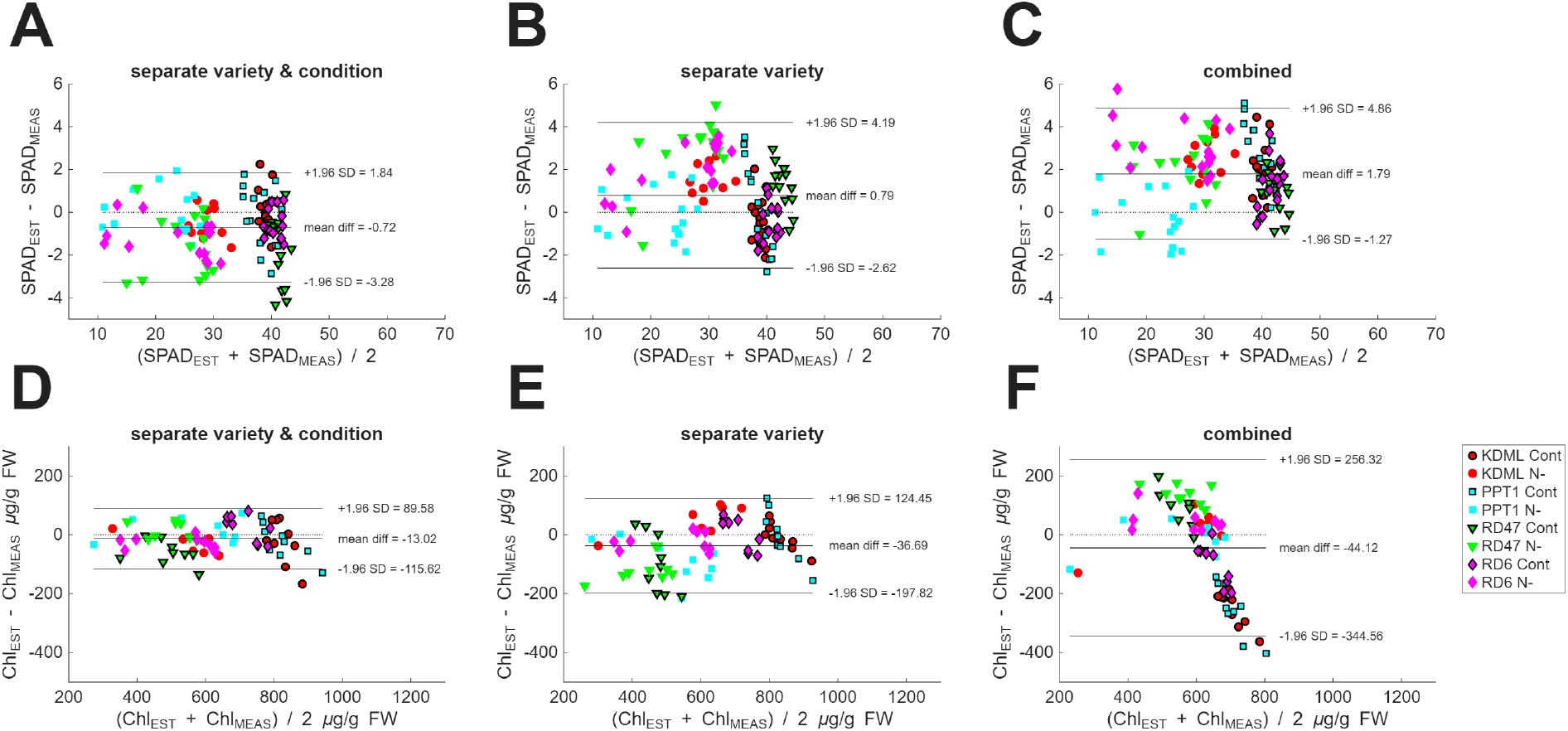
Bland-Altman plots associated with PhotoFolia image-based estimates corresponding to results presented on Fig. 6 (see text for details).

For chlorophyll estimates (Fig. 7D, E, F) both bias and LoA also increase with decreasing reference reflectance spectra specificity, and the impact of reducing specificity is more pronounced than for SPAD value. When applying separate variety and condition specific reference reflectance spectra (Fig. 7E) most chlorophyll estimates fall within +/- 100 μg/g FW of known values. However, estimates using combined reference spectra (Fig. 7F) span a range of 600 μg/g FW around the measured value. In addition, the errors are highly dependent on the chlorophyll concentration. These observations highlight the close link between leaf light transmission (SPAD) and reflection (PhotoFolia) (Xu and Ye 2023) and their less direct relationship with chlorophyll concentration.

## Discussion

This study finds that a mobile phone-based method can predict SPAD values within +/- 1.2 SPAD units (MAE, Fig. 6A) and chlorophyll concentrations within 7.2% (MAPE, Fig. 6D) of laboratory assay results. These results suggest that a mobile phone-based solution, that employs a cardboard colour reference standard, could offer a practical and cost-effective method for informing plant management decisions in the field.

### Predicting SPAD value and chlorophyll concentration from camera responses

Whereas the Beer-Lambert law implies that the logarithm of the camera response should be linearly related to SPAD values (and chlorophyll concentrations), this model did not significantly reduce prediction errors compared to a simpler linear model. Based on predicted camera responses, a purely linear model appeared to be a reasonable and practical choice. A more complex model based on the relationship between leaf spectral reflectance, camera response, and SPAD/chlorophyll values might improve prediction accuracy, but there is no clear evidence that this relationship is especially complex, and using the simplest (linear) form is unlikely to compromise practical performance.

### Accuracy of SPAD estimates

The MAE of image-based SPAD value estimates (± 1.2 units, Fig. 6A) is similar to the published accuracy of the Minolta 502+ SPAD meter (± 1.0 units). This is a reasonable expectation if the factors that are known to contribute uncertainty to image-based estimates are effectively mitigated. The variation in the relationship between measured SPAD value and PhotoFolia estimates, as represented by MAP and MAPE values (Fig. 6) might arise simply because the precise position of SPAD and PhotoFolia measurements on the leaf differed slightly. Consequently, we could not ensure that the pixel values used for image-based estimates corresponded to the same region of the leaf covered by the SPAD meter’s aperture. The measurement aperture, which is relatively small (approximately 6 mm in diameter), is ideal for measuring leaves with small areas but may, by chance, align with a leaf vein or an interveinal area, each having different light absorption properties. Although this risk was minimal for healthy rice leaves (controls), which are relatively uniform in colour, nitrogen depletion treatments caused variegation (striping) in leaf surface colour, which could easily cause the measured variations.

### Accuracy of chlorophyll estimates

The situation is more complex for chlorophyll estimates. This study does not find a consistent linear relationship between SPAD value and chlorophyll concentration, based on the simplest interpretation of the Beer Lambert law. Instead, different varieties of rice leaf grown under different conditions exhibit unique (albeit linear) relationships between SPAD value and chlorophyll concentration. Likewise, similar measured SPAD values were associated with different chlorophyll concentrations for one variety of rice grown under different conditions (RD47, Fig. 5). This implies that estimation of rice leaf chlorophyll concentration from either SPAD value or the response of a camera should factor in rice variety and growth conditions.

These results align with findings that the relationship between SPAD value and chlorophyll concentration varies across species, including birch, wheat, potato (Uddling et al. 2007), Amazonian trees (Marenco et al. 2009), rice, maize, soybean, peanut, cotton (Xiong et al. 2015), and Mexican and Pima cotton (Pauli et al. 2017).

Structural and textural features of leaves, such as ridges, hairs, and wax coatings, will influence how light interacts with the leaf, and hence the surface spectrophotometry measurements of chlorophyll concentration, but for the rice leaves used in this study the abaxial surface properties were very similar, so such surface properties are unlikely to explain the observed effects of strain and treatment. However, internal structural heterogeneity (specifically the non-uniform chlorophyll distribution between vascular bundles (veins) and mesophyll tissue) introduces a “sieve effect” that violates the homogeneity assumption of the Beer-Lambert law (Duysens 1956; Terashima 1983). Although rice exhibits parallel venation, veins consistently contain lower chlorophyll concentrations than interveinal regions. Variations in vein density, bundle sheath development, or interveinal distance between genotypes or in response to nitrogen availability could amplify measurement uncertainty. Despite these violations of the homogeneity assumption, Beer-Lambert is widely considered to provide a good approximation (Evans 1995; Evans 2003; Holloway-Phillips 2019; Terashima 1985).

It should be noted that whereas laboratory assays use mass-based chlorophyll concentration (µg/g FW) optical attenuation follows the Beer-Lambert law, which inherently gives an area-based measure of chlorophyll content (µg/cm^2^) - the total pigment encountered along the optical path through the leaf. Consequently, variations in leaf thickness, specific leaf weight, or internal structure that affect optical path length will confound attempts to estimate mass-based chlorophyll using a generic optical model. However, the present study has an agronomic objective: to demonstrate that a low-cost optical method can replicate laboratory assay results, which conventionally report chlorophyll on a mass basis (µg/g FW). This metric aligns with tissue testing standards used by farmers and agronomists to assess nitrogen status and guide fertilizer decisions (Li 2022). While the SPAD meter is effective for estimating area-based chlorophyll content, it performs poorly when inferring mass-based concentration without correction for specific leaf weight (Li 2011).

The key innovation of our approach is the use of cultivar- and condition-specific reference spectra. This specificity enables ColourWorker to achieve laboratory-comparable accuracy for mass-based chlorophyll estimation. This is evidenced when comparing Figures 6C and 6F: a generic model predicting SPAD values (an area-based proxy, Figure 6C) performs well across diverse samples, whereas a generic model predicting mass-based chlorophyll concentration (Figure 6F) performs poorly. This disparity arises because optical signals inherently respond to area-based pigment content; without accounting for structural variation, mass-based estimation is confounded.

However, when reference spectra are tailored to specific cultivars grown under defined conditions, the model implicitly learns the characteristic relationship between the optical signal and mass-based chlorophyll for that genotype. Within a given rice cultivar, leaf thickness and internal structure tend to be relatively uniform. Under these constraints, the primary source of variation in light absorption becomes the chlorophyll density/distribution within the leaf tissue (µg/g FW), allowing accurate mass-based inference despite the fundamental physics favouring area-based units. For completeness, we repeated our analysis using area-based chlorophyll units (µg/cm^2^), which show stronger alignment with SPAD-type measurements and are provided in Supplementary Material (Figures 1-3).

In addition, chlorophyll interactions with incident and scattered photons may be affected by the maturity of chloroplasts and or how these are oriented in the leaf with respect to the light path (Howard et al. 2019; Kong and Wada 2014; Xiong et al. 2015). Consequently, the complex internal structure of leaves, and the fact that chloroplasts dynamically change position in response to the light environment and growth conditions (Xiong et al. 2015) may explain why we observed different relationships between plants grown under control and nitrogen depleted conditions. For example, nitrogen starvation can reduce chlorophyll a, and accessory pigments such as carotenoids (Cetner et al. 2017) and also the leaf thickness due to reduced number of cell layers (Yamamoto et al. 2002). Despite making measurements at similar times during the day, this suggests that genetic factors (variety dependence of nitrogen assimilation) as well and nutrient availability (treatment dependence) could both be relevant.

This study made no attempt to investigate causes of these variations, but while estimates of SPAD value from leaf images were relatively insensitive to the specific choice of reference reflectance spectra when using PhotoFolia, the reference spectra used substantially impacted prediction of chlorophyll concentration. Unsurprisingly, a generic linear model for predicting chlorophyll concentration that combined variety- and condition-specific reflectance spectra was ineffective (Fig. 6F).

We note also that although the spectrophotometrically measured chlorophyll concentration should be a ground truth value, solvent-based extractive techniques produce several sources of uncertainty. While every attempt was made to control laboratory factors, such as solvent quality, grinding technique, post sampling storage, extraction time & temperature, chlorophyll is a biologically active molecule, a potentially unstable pigment, which can undergo photo-degeneration during sample preparation and measurement (Petrović et al. 2017). In addition, the absorption-based measurements recovered from the chlorophyll extract are themselves prone to a variety of uncertainties ranging from spectrometer calibration errors at critical wavelengths to interference from additional pigments (Aminot and Ray 2000; Riemann 1978). Also, as in the case of the SPAD value measurements, the chlorophyll extraction was not made from the same identifiable region of the leaf that was photographed when making image-based estimates.

It is difficult to quantify the contributions of these different sources of uncertainty. And despite ostensibly being a ground truth measurement, spectrophotometry is likely to exhibit a greater level of uncertainty compared to the raw measurements recovered from hardware SPAD meters and camera images (Tan et al. 2021), but crucially, assay-based spectrophotometry is much less affected by the way that chlorophyll is distributed in the leaf. So ultimately, it is the appropriate ground truth for quantifying the true chlorophyll content.

### Practical utility of image-based leaf chlorophyll measurement

This study shows that image-based measurements of leaf chlorophyll implemented by PhotoFolia has similar performance and limitations to SPAD for informing agricultural practise. This study evaluated PhotoFolia using several models of mobile phone under normal (diffuse) natural daylight illumination and demonstrated that the method is robust to differences in camera hardware and to the typical variation in ambient lighting encountered in field conditions. However, the effect of controlled or extreme illumination variation - such as changes in intensity, angle, or spectral distribution - was not systematically investigated here. Nor was any attempt made to systematically compare performance across a wide range of mobile devices. No attempt was made to regulate or standardise the spectral properties of the lighting during image acquisition; instead, images were captured under whatever ambient daylight was present at the time, reflecting real-world use.

Converting an optical signal to absolute chlorophyll concentration necessitates a calibration model, a constraint applicable to all optical methods including the SPAD meter. PhotoFolia addresses this by utilising a pre-defined calibration based on reference reflectance spectra and ground-truth data (Fig. 2). The distinction lies in the implementation: PhotoFolia integrates this calibration data into the app, requiring only a selection of leaf type and condition from the user. Conversely, instruments like the Minolta SPAD 502+ require the user to separately correlate index readings with chlorophyll concentrations. PhotoFolia maintains scientific rigor regarding the calibration process while significantly reducing the procedural complexity for the end-user.

Further, an image-based chlorophyll measurement procedure that can be implemented on widely available mobile phones is obviously convenient. Moreover, mobile phones provide GPS and are often operated on-line, enabling integration of geo-located chlorophyll concentration data with cloud-based analytics.

However, PhotoFolia is not an entirely hardware independent solution, it still requires a colour standard to enable a calibrated camera response. Some care is needed during image acquisition to ensure consistent lighting across the colour standard and target leaf, and to avoid occlusion, shadowing, or uneven illumination of the colour standard. However, a key advantage of this approach is that it captures a visible image of the plant and colour standard, which the user can immediately review on their device. This allows the user to make an informed judgment about image quality - such as checking for proper framing, focus, and lighting - before proceeding with analysis. This is not possible with traditional “blind” handheld devices that provide a measurement without capturing or displaying a visual record. Many potential sources of uncertainty in the final chlorophyll concentration estimate can be mitigated. In addition, the use of colour cards is familiar to some potential users.

The main uncertainty at this point is precisely how information about chlorophyll concentration—or indeed any plant trait inferred from spectral reflectance or imagery—can translate into actionable insights for farmers. What growers ultimately seek is reliable, real-time guidance on leaf nitrogen status, which directly informs fertilizer decisions and correlates with crop protein content and yield potential. While chlorophyll is often used as a proxy for nitrogen, this relationship can vary across genotypes, growth stages, and environmental conditions. Nevertheless, the fact that PhotoFolia enables chlorophyll concentration estimates from leaf images that are within 10% (MAPE, Fig. 6D) of a laboratory-based assay—almost instantly and at virtually no cost—suggests tangible potential to support on-farm decision-making, particularly if coupled with calibration strategies that link chlorophyll to nitrogen under specific agronomic contexts.

## Acknowledgements

J. Anderson contributed to the development of analytical and data processing methods, data analysis, software development, and manuscript. K. Boonyaves and H. Okamoto contributed to the experiment design, data collection, and to the writing of this manuscript. N. Karaket, W. Chuekon, and N. Chabang provided valuable support in gathering and organising the experimental data. M. Garvie developed the mobile phone application used in the study. Daniel Osorio and K. Supaibulwatana contributed to the planning the study, experimental design and preparation of the manuscript. The authors thank the anonymous reviewers for their constructive comments and insightful suggestions, which improved the quality and clarity of this manuscript.

This research was supported by the Mahidol-Sussex seed fund.

