## Supplementary material for "Image-based leaf SPAD value and chlorophyll measurement using a mobile phone: enabling accessible and sustainable crop management"

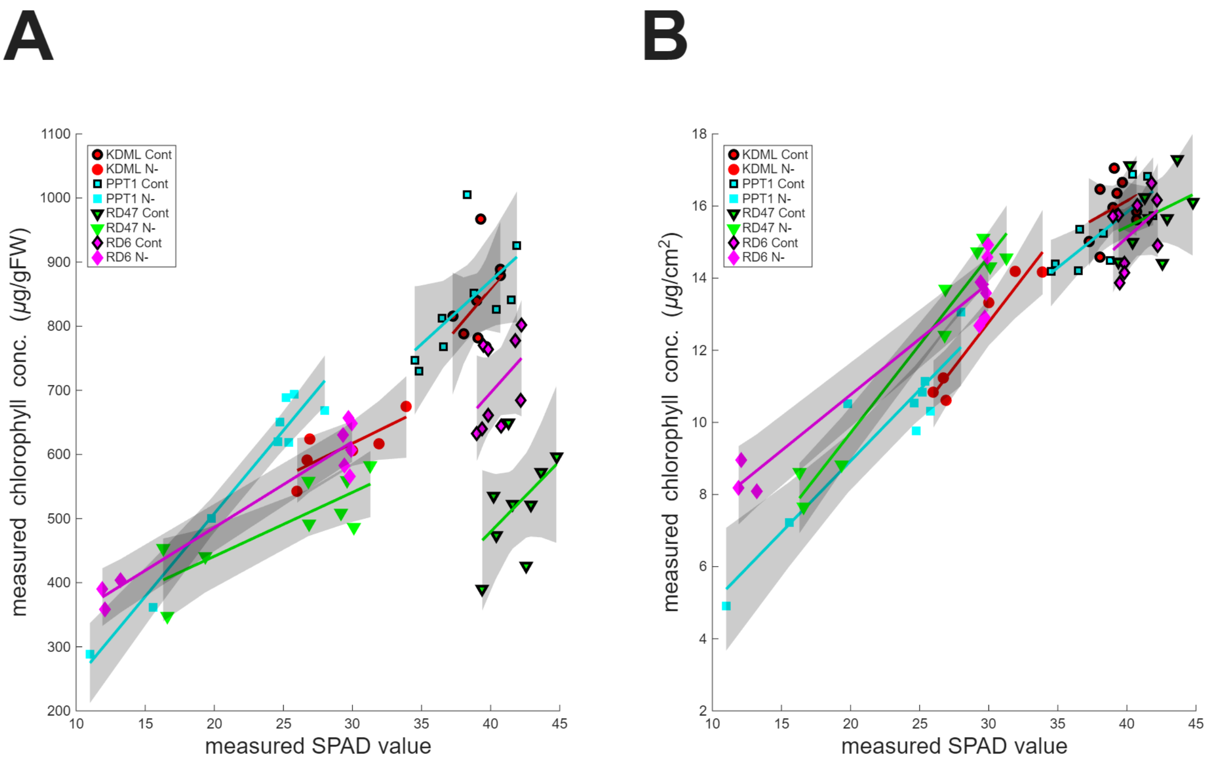


**Figure 1. Relationship between chlorophyll content and SPAD measurements across rice varieties and growth conditions. (A) Mass-based chlorophyll concentration ($\mu g/g$ FW) versus SPAD value, showing substantial scatter due to variety- and condition-specific differences in leaf structure (reproduction of Figure 5 in main paper for comparison). (B) Area-based chlorophyll content ($\mu g/cm^2$) versus SPAD value. Area-based values were derived from mass-based measurements by normalising to leaf area (eqn. 1), where leaf fresh weight was recorded prior to chlorophyll extraction and leaf area was estimated from digital images. Leaf length was fixed at 3 cm; width was determined from the fraction of the colour standard aperture (10 mm diameter) covered by the leaf in each image. The tighter correlation in the right panel reflects the shared physical basis of area-based chlorophyll content and SPAD measurements: both respond to total pigment encountered along the optical path (Beer–Lambert law). In contrast, mass-based concentration is confounded by variation in leaf mass per area (LMA), as structural differences alter optical path length without changing pigment per unit mass.**

$$\mathrm{Chl\_area}\ (\mu g/cm^2) = \mathrm{Chl\_mass}\ (\mu g/g\ \mathrm{FW}) \times \mathrm{Fresh\ Weight}\ (g) / \mathrm{Leaf\ Area}\ (cm^2) \qquad \text{(eqn. 1)}$$

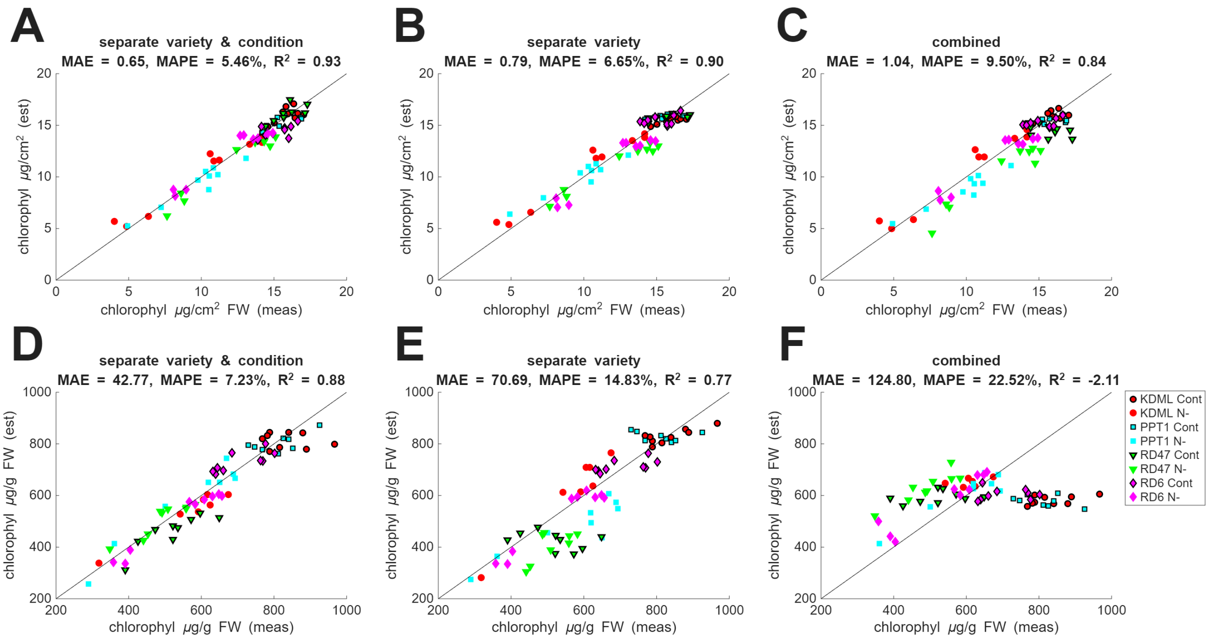


**Figure 2. Comparison of image-based chlorophyll estimates (PhotFolia app) against laboratory measurements for area-based content (panels A-C) and mass-based concentration (panels D-F), illustrating the impact of reference spectrum specificity. Seven-week-old leaves from four rice varieties (KDML105, PPT1, RD47, RD6) grown under two nitrogen regimes, control (2.056 mM) and nitrogen-deficient (0.26 mM) - were analysed. (A, D) Separate variety & condition reference spectra; (B, E) variety-specific spectra with conditions merged (C, F) a single generic spectrum for all samples. Across equivalent specificity levels, area-based estimates consistently outperform mass-based estimates (compare A vs. D, B vs. E, and C vs. F), as evidenced by tighter agreement with laboratory measurements. This superior performance arises because area-based chlorophyll content (µg/cm²) aligns with the optical physics of light attenuation (Beer–Lambert law), whereas mass-based concentration (µg/g FW) is confounded by variation in leaf mass per area (LMA) - i.e., structural differences in leaf thickness and density that alter optical path length without changing pigment per unit mass (see Discussion).**

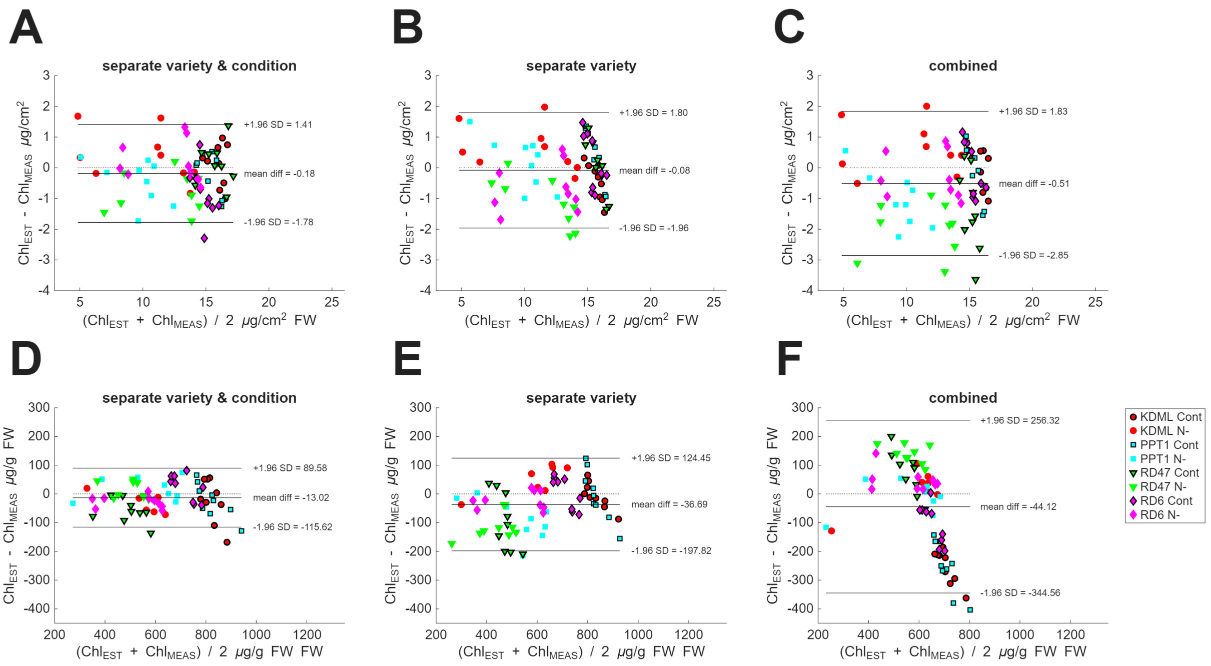

**Fig. 3. Bland–Altman plots comparing PhotoFolia image-based estimates against laboratory measurements for area-based chlorophyll content (top row) and mass-based concentration (bottom row), corresponding to the accuracy assessments in Supplementary Material Fig. 2. Panels show results using reference spectra of decreasing specificity. Merging variety and condition specific populations, panels B and C, introduces systematic error correlations, with bias increasing at high chlorophyll concentrations. This pattern may arise because nitrogen-sufficient and nitrogen-depleted leaves exhibit different reflectance - chlorophyll relationships due to treatment-induced changes in leaf anatomy (e.g., thickness, mesophyll structure) and pigment composition, even at equivalent chlorophyll content. A single linear model fitted across both populations may represent a compromise. This is corroborated in Supplementary Material Fig. 1, where separate linear fits for each treatment group show steeper slopes for nitrogen-depleted leaves than controls at high chlorophyll concentrations. The emergence of structure in the residuals upon merging reference libraries underscores the importance of condition-specific calibration for accurate estimation from optical measurements in this context but may also reflect the choice of a simple linear model when summarising the overall relationship.**
